# Entraining corticocortical plasticity changes oscillatory activity in action control and inhibition

**DOI:** 10.1101/2020.06.10.142398

**Authors:** Alejandra Sel, Lennart Verhagen, Katharina Angerer, Raluca David, Miriam Klein-Flügge, Matthew Rushworth

**Affiliations:** Wellcome Centre for Integrative Neuroimaging (WIN), Department of Experimental Psychology, University of Oxford, Oxford, OX1 3UD, UK; Centre for Brain Science, Department of Psychology, University of Essex, Wivenhoe Park, Colchester, CO4 3SQ; Donders Institute for Brain, Cognition and Behaviour, Radboud University Nijmegen, Nijmegen, 6525 HR, the Netherlands

**Keywords:** action control, inhibition, cortico-cortical repetitive paired associative transcranial stimulation, paired-pulse TMS, ventral pre-motor cortex, primary motor cortex, Hebbian plasticity, EEG, beta oscillations, theta oscillations

## Abstract

Oscillatory activity may reflect interactions between brain areas[1]. Here we tested whether inducing corticocortical plasticity in a specific set of connections changes oscillatory activity and cortico-cortical interactions and, if this is the case, whether the changes manifest in a manner that is behaviour state-dependent. We either increased or decreased the influence of activity in human ventral premotor cortex (PMv) over activity in primary motor cortex (M1) using cortico-cortical paired associative stimulation (ccPAS)[2, 3]. Before and after stimulation participants performed a Go/No-Go task. While M1 TMS pulses revealed the excitatory state of the motor system at specific time points, the electroencephalogram (EEG) revealed the evolution of oscillatory activity dynamics in the motor system over several hundreds of milliseconds before, during, and after each movement. Augmenting cortical connectivity between PMv and M1, by evoking synchronous pre- and postsynaptic activity in the PMv-M1 pathways, led to a state-dependent modulation of the causal influence of PMv over M1, and at the same time, enhanced oscillatory beta and theta rhythms in Go and No-Go trials, respectively. No changes were observed in the alpha rhythm. The plasticity induction effect was dependent on PMv-M1 stimulation order; the opposite patterns of results were observed after an equal amount of stimulation of PMv and M1 but applied in a temporal pattern that did not augment PMv’s influence over M1. These results are consistent with Hebbian principles of synaptic plasticity[4] and show that artificial manipulation of cortico-cortical connectivity produces state-dependent functional changes in the spectral fingerprints of the motor circuit.

## Results

Across four experiments we examined the possibility of selective potentiation of physiological connectivity in the PMv-M1 corticocortical pathway. We recorded motor-evoked potentials (MEPs) from the left first dorsal interosseous (FDI) muscle – Experiments 1+2, and EEG oscillatory activity – Experiments 3+4, while participants performed a Go/No-Go task in two blocks (Supplemental Material). Between the task blocks, we used a particular type of TMS –ccPAS, as our experimental manipulation, comprising repeated paired stimulation of PMv and M1. In Experiments 1 and 3, we applied 15 minutes of ccPAS over PMv followed by M1 (PMv-M1-ccPAS; each PMv pulse was followed by an M1 pulse at an 8ms inter-pulse interval, IPI) between the two Go/No-Go task blocks (referred to as Baseline and Expression blocks). Experiments 2 and 4 investigated whether such changes in PMv-M1 connectivity were dependent on ccPAS stimulation order by reversing the order of ccPAS stimulation, i.e. applying the first paired pulse over M1 and the second pulse over PMv (Fig.1). Exactly the same number of pulses were applied to PMv and M1 in all experiments, with the same IPI. All the experimental procedures were approved by the Medical Science Interdivisional Research Ethics Committee (Oxford RECC, No. R29477/RE004).

**Figure 1:**
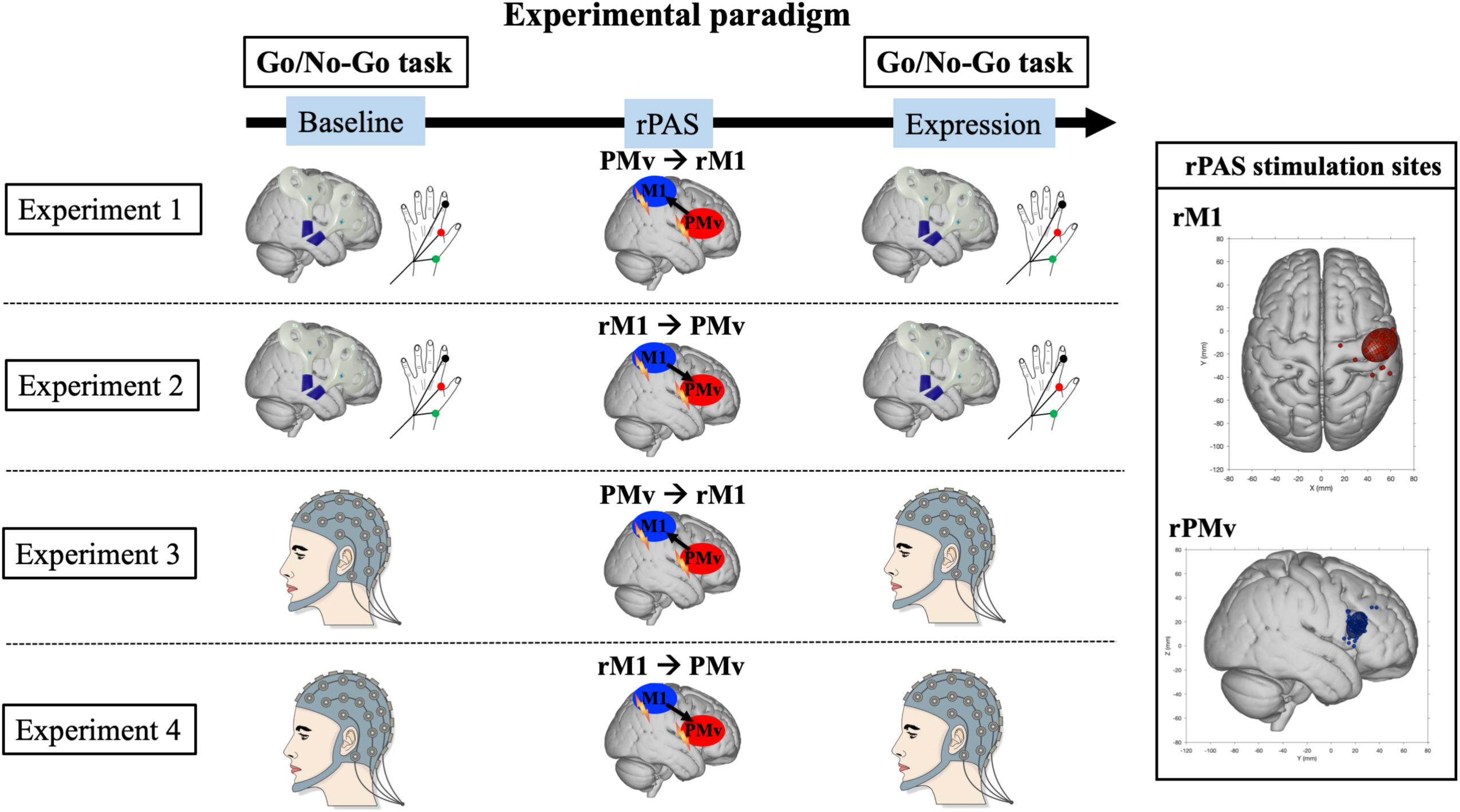
**Left:** Experimental design and setup for all experiments. The corticocortical repetitive paired associative stimulation (ccPAS) period was preceded (Baseline) and followed (Expression) by Go/No-Go task blocks. TMS-induced motor evoked potentials (MEPs) – Experiment 1 and 2, and EEG motor responses – Experiment 3 and 4, were recorded during the task blocks. **Right:** Individual subject scalp hotspot (filled circles) and 95% group confidence ellipses for rM1(red) and rPMv (blue) locations for all experiments in standardized MNI space.

### TMS Experiments

In Experiments 1 (n=18) and 2 (n=17), we first characterised the causal relationship between PMv and M1 by testing the influence of PMv over M1 during the Go/No-Go task, and examining whether this influence was state-dependent (i.e. different for Go *vs* No-Go trials) by contrasting MEP amplitudes across conditions. The MEPs provided our dependent measure; we applied either single-pulse TMS (spTMS) over right M1 (125ms after stimulus onset) [5, 6], or paired-pulse TMS (ppTMS) over right PMv (conditioning pulse) and right M1 (8ms IPI) [2, 3, 7, 8] and recorded motor-evoked potentials (MEPs) from the left first dorsal interosseus (FDI) muscle while participants performed the Go/No-Go task before and after ccPAS. Stimulation of PMv and adjacent frontal cortex has been shown to alter activity in M1 in previous experiments but the nature of the influence (facilitatory or inhibitory) depends on the task, trial context, and timing [5-8]. As reported previously [3, 5-9], before ccPAS (Baseline) MEPs were larger when M1 TMS was preceded by a conditioning pulse over PMv on Go trials (Fig.2A/2B (left)-light *vs* dark blue bars) suggesting that PMv exerted an excitatory influence over M1 in both Experiments 1+2 (t(11)=-2.235, p=0.047, d=0.649; statistical test conducted across the paired datasets from the twelve individuals who participated in Experiments 1+2 in a counterbalanced order).

**Figure 2:**
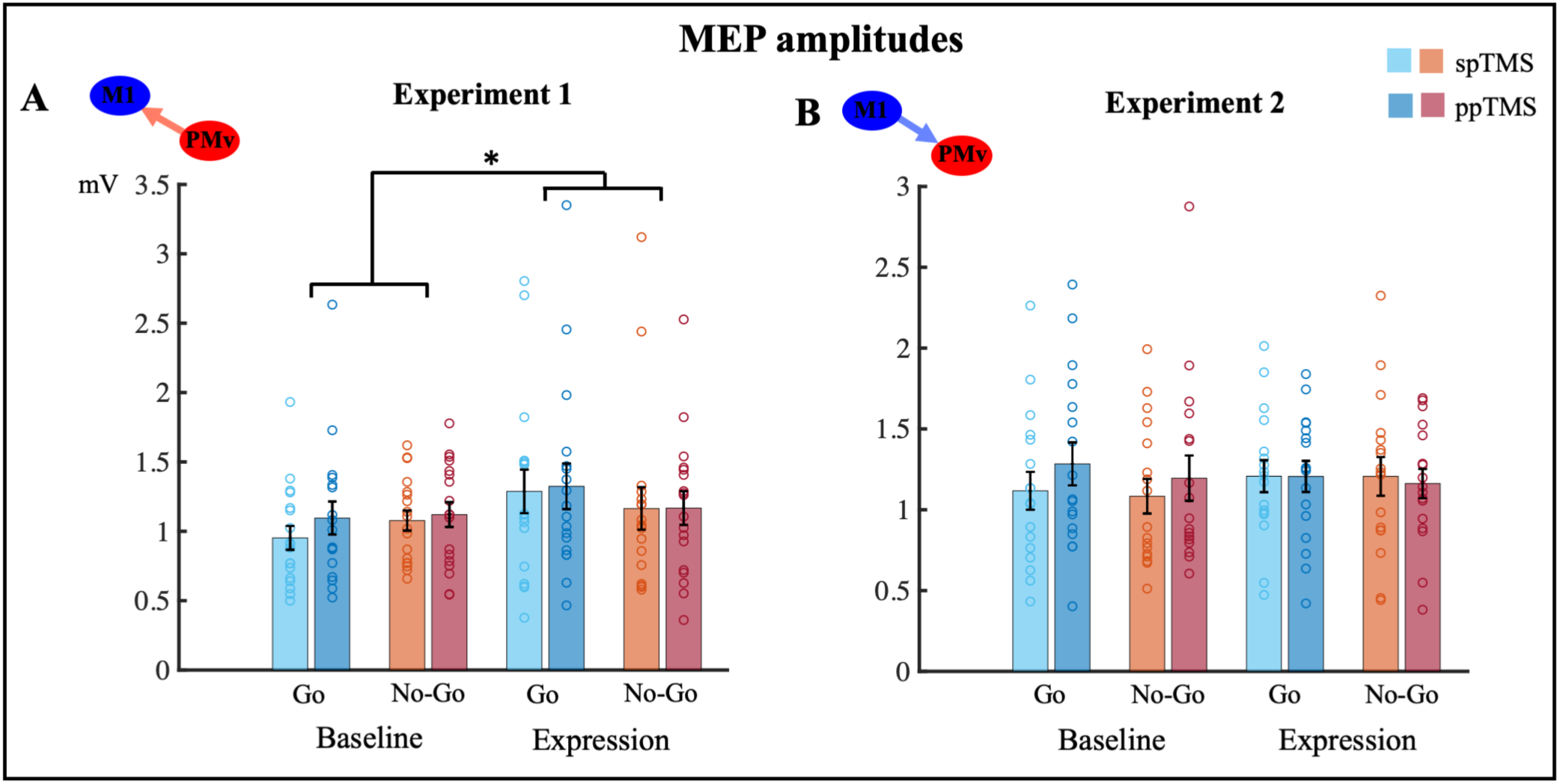
Group mean MEP amplitudes measured with either spTMS over M1 or ppTMS over PMv/M, in Go and No-Go trials, recorded before (Baseline) and after (Expression) ccPAS in Experiment 1 (left) and Experiment 2 (right). Error bars represent SEM, single dots represent individual data points

To examine potentiation of physiological connectivity in the PMv-M1 corticocortical pathway [2, 3], we contrasted the neural excitability of M1 by means of MEPs recorded during the Go/No-Go task before and after each type of ccPAS. The analysis of MEP amplitudes in Experiment 1 revealed a significant interaction between block (Baseline, Expression) and trial type (Go, No-Go) (*F*(1,17)=6.74, *p*=.019, η_p_^2^ =.28). Subsequent analysis demonstrated that the interaction was due to the impact that ccPAS had on Go trials as opposed to No-Go trials. When comparing the MEPs recorded for each trial type in Expression and Baseline blocks (computed as Expression Block MEP–Baseline Block MEP), the change in MEP size was greater on Go than No-Go trials (*t*(17)=2.60, *p*=.019, d =0.6136); (Fig.2). Crucially, PMv-M1 ccPAS increased M1 activity on Go trials (Fig.2A(left)-Exp1; blue bars are higher on right than left); repeated stimulation of PMv followed by M1 leads to an increased excitability in M1 during Go trials. By contrast, in Experiment 2, the reverse order of ccPAS stimulation (M1-PMv) did not modulate MEP amplitudes in either Go or No-Go trials (all *p*>0.05).

These results demonstrate two types of influence exerted by PMv over M1. First, PMv exerted a transient influence on M1 each time a pulse was applied to PMv. At Baseline in both Experiments 1+2, conditioning pulses of PMv enhanced the MEPs induced by M1 TMS on Go trials when neurophysiological studies show that PMv neural activity is inducing changes in M1 activity [10-12]. Second, PMv-M1 ccPAS in Experiment 1 caused sustained changes in activity in M1 whenever a movement was subsequently made (when PMv exerts an influence over M1 [10-12]) regardless of whether or not a further conditioning pulse was applied. The sustained change was present on Go trials in contrast to No-Go trials. There was no evidence of a similar sustained change in M1 when an equal number of TMS pulses were applied to M1 and PMv but in the opposite order during M1-PMv ccPAS (Experiment 2).

### TMS-EEG Experiment

In Experiment 3 (N=16) and 4 (N=17), we investigated whether Hebbian-like physiological changes in the PMv-M1 corticocortical pathway lead to modulation of both fast (transient) and slow (sustained) EEG oscillatory dynamics associated with action control. We contrasted the effects of ccPAS on time-frequency oscillatory responses, i.e. the ‘cortical entrainment effect’ (computed as Expression–Baseline block for Go and No-Go trials, separately) between the two groups of participants. We focused on motor-relevant frequency bands theta, alpha and beta (4-30Hz) in mid-frontal electrodes previously associated with top-down executive control [13-18].

The results of a cluster-based permutation analysis (Supplemental information) revealed that ccPAS had a significant impact on motor-related beta and theta bands. No ccPAS effects were found in the alpha band. That is, the cortical entrained effects significantly differed for Go and No-Go trials between the two participant groups tested before and after the two varieties of ccPAS in Experiments 3+4 in the beta band (19–24Hz; *p*=0.018), as well as in the theta band (4–10Hz; *p*=0.008;) between 0.15 and 1.2s after the stimulus onset in fronto-central sites. Following these results, we contrasted the cortical entrained effect across the two participant groups for Go and No-Go trials, separately. In the beta band, post-hoc analyses showed that the PMv-M1 ccPAS led to an increase of beta synchronization (0.7-1.2s after stimuli onset, consistent with the time of the post-movement beta rebound, PMBR) in the Expression *vs* the Baseline block only for Go trials, whereas the opposite effects were found in Go trials when the ccPAS order was reversed (p=0.002–Fig.3A-left**). No significant differences in the beta band were observed for No-Go trials (p>0.05) (Fig.3A-right). The PMBR is typically observed over motor and premotor areas following the completion of a movement [13, 14]. Beyond the traditional notion of the PMBR as a short-lasting state of motor deactivation, or a “resetting” of the underlying neural networks [19], several potential roles for this PMBR have been suggested. Accumulating evidence suggests that the PMBR reflects a short-lasting state of active inhibition of motor cortex networks after movement completion [13, 20, 21]. It is possible that the increased PMBR observed in Go trials during Expression reflects an augmentation of active inhibition from PMv over M1 following strengthening of PMv’s influence over M1 through ccPAS. The projections from PMv to M1 are excitatory but many of these projections are onto inhibitory interneurons in M1 [22]. Thus, PMv exerts both inhibitory and facilitatory influences over M1 and both of these influences can be augmented by PMv-M1-ccPAS. Moreover, the observation of the opposite effects on beta synchronization on Go trials when reversing the order of the ccPAS stimulation in Experiment 4 are in line with previous evidence showing contrasting effects of reversed *vs* forward order ccPAS on M1 cortical excitability, as well as on functional connectivity in motor networks [2, 3].

**Figure 3:**
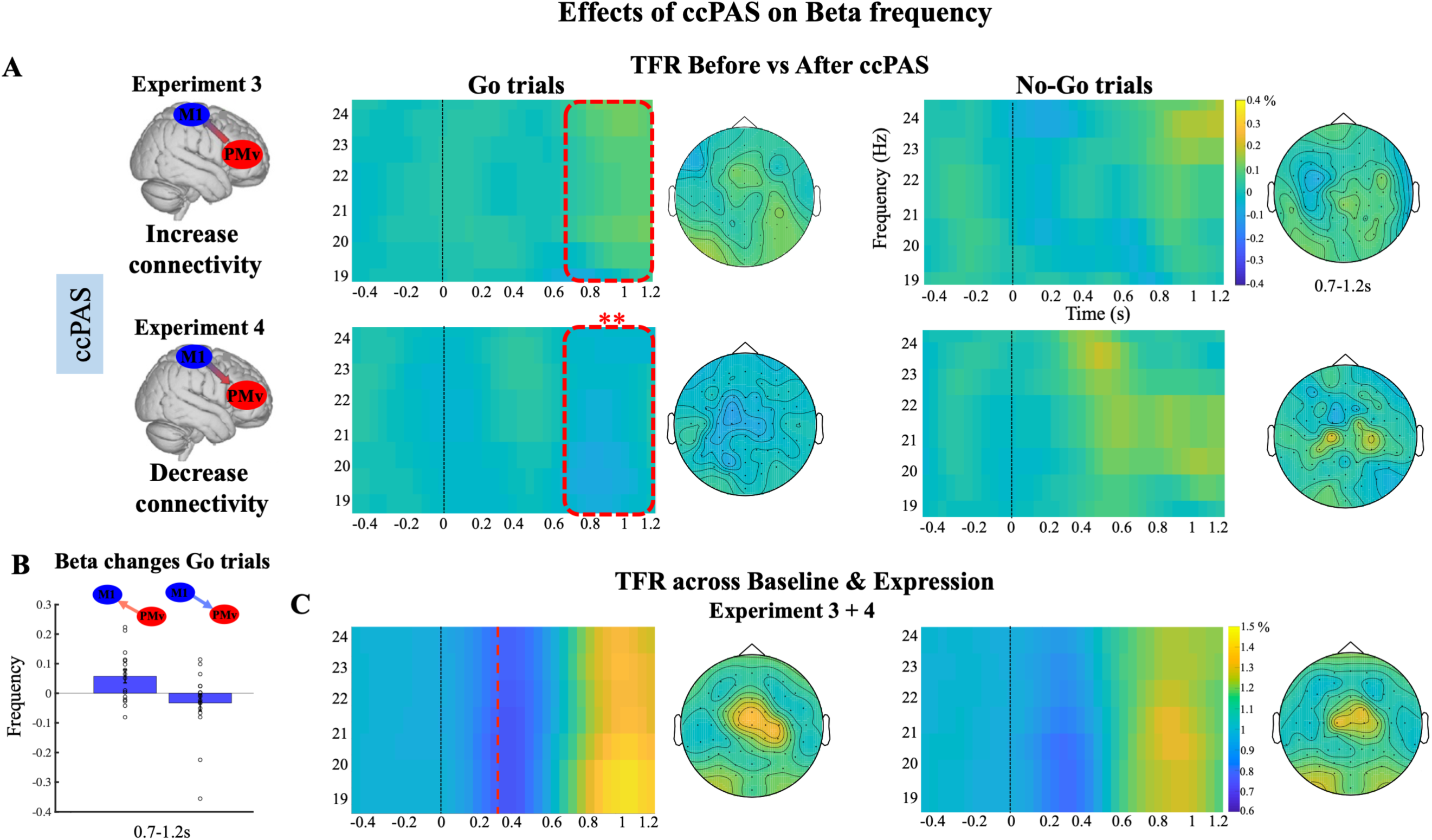
**A and C:** EEG time-frequency responses (TFR) in the beta band (19-24Hz) in fronto-central sites (negative cluster; C4, CZ, FC2, CP2, FCZ, C1, C2, FC4, CP4, CPz) time-locked to the onset of the Go/No-Go stimuli, computed as **(A)** the difference between Baseline and Expression blocks, **(C)** the mean of Baseline and Expression blocks collapsing across Experiment3+4. While **C** shows the PMBR effect was especially prominent in the Go trials, **A** illustrates how this changed as a function of the two types of ccPAS used in Experiments 3 and 4. Dashed red square in **A** indicates the time window (0.7-1.2s) where significant modulation in beta responses after ccPAS were found. Dashed red line in **C** indicates the mean RT across Baseline and Expression for Go trials in both participant groups (mean= 352.36s). **B:** mean beta frequency increase (PMv – M1 ccPAS) and decrease (M1 - PMv ccPAS) computed as the difference between Expression and Baseline, in Go trials, in the 0.7-1.2s time window. Error bars represent SEM, single dots represent individual data points

In No-Go trials, PMv-M1 ccPAS led to a significant increase in theta power, whereas theta power decreased after reversed order M1-PMv ccPAS (p=0.002; 0.15-1.2s after stimuli onset) (Fig.4). Several findings have linked increased theta power in mid-frontal regions to top-down executive control and action reprograming during response conflict and motor inhibition, e.g. after a No-Go command [15-18]. Notably, theta oscillatory changes increase with the level of response conflict reflecting a larger top-down influence over motor circuits [17, 18]. Therefore, the increased theta power in No-Go trials after PMv-M1 ccPAS observed in Experiment 3 suggests augmented top-down motor control in response conflict, whereas the reversed order M1-PMv ccPAS suggests decreased executive control over motor output in No-Go trials. No ccPAS effects on theta power were found in Go trials (p>0.05) (Fig.4).

**Figure 4:**
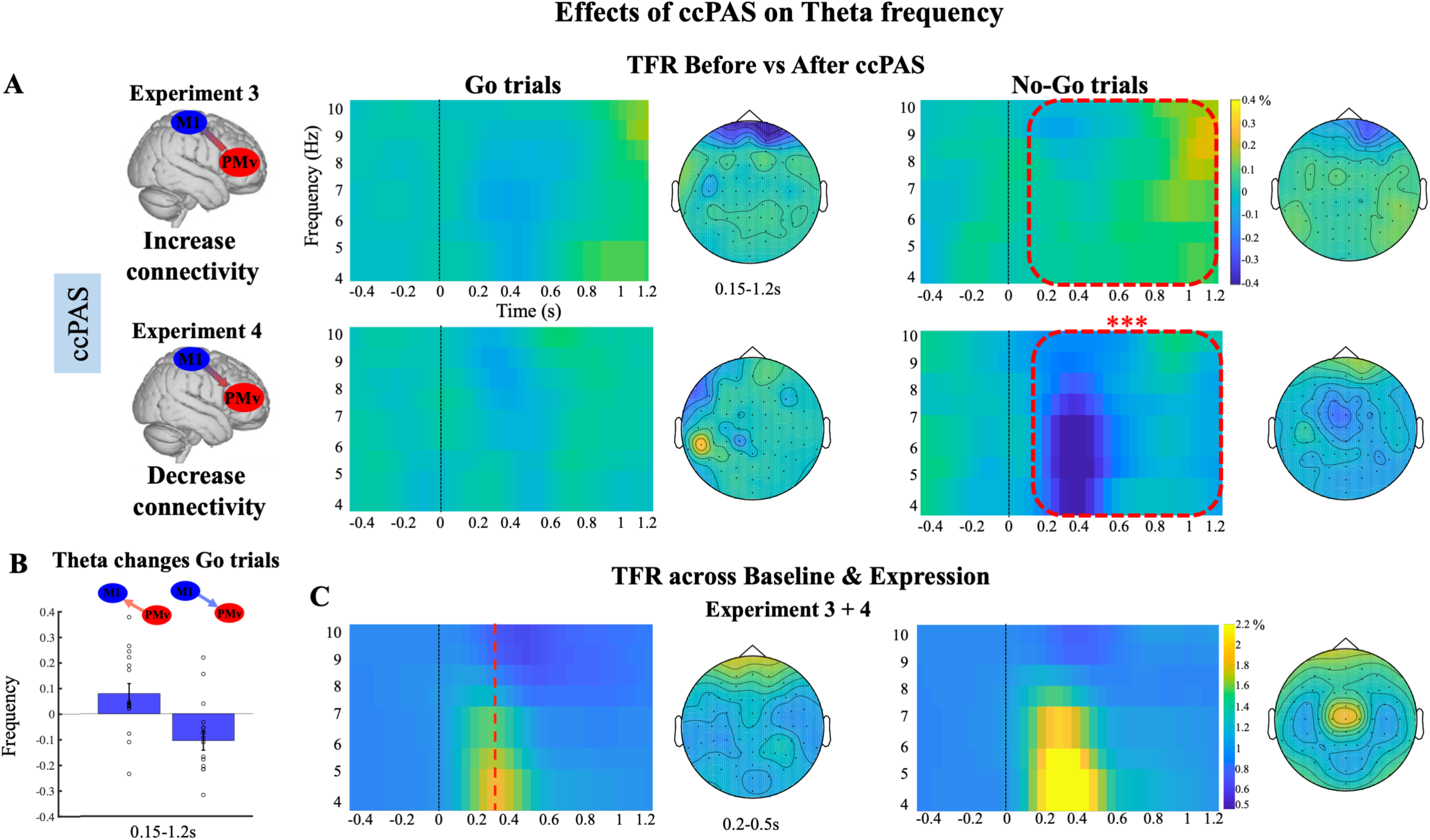
**A and C:** EEG time-frequency responses in the theta band (4-10Hz) in fronto-central sites (positive cluster; C3, C4, CZ, FC1, FC2, FCZ, C1, C2, FC3, FC4, CP4, CPZ) time-locked to the onset of the Go/No-Go stimuli, computed as **(A)** the difference between Baseline and Expression blocks, **(C)** the mean of Baseline and Expression blocks collapsing across Experiment 3+4. While **C** shows the theta effect that was especially prominent in the No-Go trials, **A** illustrates how this changed as a function of the two types of ccPAS used in Experiments 3 and 4. Dashed red square in **A** indicates the time window (0.15-1.2s) where a significant modulation in theta responses after ccPAS was found. Dashed red line **in C** indicates the mean RT across Baseline and Expression for Go trials in both participant groups (mean= 352.36s). **B:** Mean theta frequency increase (PMv – M1 ccPAS) and decrease (M1 - PMv ccPAS) computed as the difference between Expression and Baseline, in No-Go trials, in the 0.15-1.2s time window. Error bars represent SEM, single dots represent individual data points

Oscillatory signals can reflect both transient, evoked activity and sustained, induced neural oscillations. Evoked responses are phase-locked to external stimuli, whereas induced oscillations are not. It is possible that beta and theta modulations occuring after ccPAS reflect changes in either one or other neurophysiological mechanism or even a mixture of both mechanisms. In order to understand the nature of the ccPAS modulations we carried out an analysis to identify any evoked oscillatory effects by computing the phase coherence across trials (inter-trial linear coherence–ITLC) for each condition. First, we determined which parts of the Go/No-Go cue-related activity were evoked or sustained regardless of ccPAS. We observed phase coherence across all frequencies tested (4-30Hz) from 0.15 to 1.2s after stimulus onset but this was particularly obvious in the theta range during an early short-lived period around 0.3s after stimulus presentation (Fig.S3–yellow area in all four right hand panels). In comparison to Go trials, No-Go trials were associated with stronger, transient, evoked activity in the theta band accompanied by milder sustained changes in theta, alpha, and beta activity. Next we examined the impact of ccPAS; while ccPAS modulated the amplitude of both early, evoked components and sustained changes of the theta oscillations in No-Go trials (Fig.4A(right)–dashed red area) and sustained changes in beta oscillations in Go trials (Fig.3A(left)– dashed red area), it did not modulate the phase consistency either in the theta or the beta band (Fig.S3, comparable phase coherence between Baseline and Expression, before and after ccPAS, for Go/No-Go trials).

## Discussion

The current findings demonstrate that it is possible to entrain oscillatory activity in specific frequency bands in a manner that is specific to behavioural context (Go or No-Go trials) by strengthening or weakening specific cortico-cortical pathways using ccPAS. We show that PMv-M1 ccPAS, which by stimulating PMv neurons immediately prior to M1 neurons, evokes synchronous pre- and postsynaptic activity in the PMv-to-M1 pathway, leads to a state-dependent augmentation of PMv’s influence over M1. This was apparent in increased MEPs, indexing excitability of motor circuits, in Experiment 1 after PMv-M1 ccPAS. The MEPs elicited by the application of single pulses of TMS to M1 on Go trials were larger, regardless of whether or not they were preceded by conditioning pulses applied over PMv. Experiments 3+4 demonstrate that ccPAS leads to state-dependent changes in beta and theta oscillatory activity. PMv-M1 ccPAS led to increased beta power in the PMBR in Go trials. Decreases and increases in beta frequency oscillations have, respectively, been linked to action initiation and cessation [1, 23]. In addition PMv-M1 ccPAS led to increased theta power when there was greater demand for motor control in No-Go trials [24]. By contrast, the opposite beta and theta patterns were seen after reversed order M1-PMv stimulation, which is unlikely to lead to simultaneous pre- and post-synaptic activity in the PMv-M1 pathway. These results demonstrate that it is possible to entrain the cortical oscillatory dynamics of action control by repeated stimulation of a directed projection in a specific motor circuit.

The current results accord with Hebbian principles of spike-timing-dependent plasticity [4], whereby the firing of presynaptic PMv cells before the postsynaptic M1 cells leads to long-term potentiation whereas the firing of postsynaptic activity before presynaptic activity usually induces long-term depression. They also accord with previous demonstrations that reversed *vs* forward order ccPAS exerts opposite effects on M1 cortical excitability and functional connectivity within motor networks [2, 3].

Notably, these observations imply that transmission of causal influences between PMv and M1 is linked to state-dependent channels of communication tuned to specific frequencies. Here, the beta rhythm for action initiation and cessation on Go trials and the theta rhythm for action inhibition on No-Go trials. The route between right PMv and adjacent inferior frontal cortex and-M1 has been linked to both action initiation and inhibition [3, 5, 9, 25, 26]. The findings demonstrate that changes in physiological connectivity between two brain areas manifest in altered oscillatory activity patterns. Such results also imply that different cortical rhythms in the beta and theta range are associated with distinct functional roles in motor control and inhibition[25].

PMv-M1 cortico-cortical ccPAS selectively modulated induced beta oscillatory activity at the time of movement completion (there was no evidence for stimulus-locked evoked beta responses). This result shows that PMv exerts its influence over M1 in a functionally specific manner, suggesting a distinctive contribution to active motor inhibition after movement completion associated with resonant activity in the beta range [14]. Such a pattern is consistent with the suggestion that a right inferior frontal region exerts inhibitory control over M1 via PMv [5, 26-28, 29]. These findings not only extend our understanding of ccPAS plasticity induction [2, 3], they reinforce hierarchical models of action control supporting the powerful effect of PMv over M1 output [29, 30] and highlight the importance of beta oscillations in the feedforward influence of premotor cortex on motor cortex [13, 31].

In contrast to the PMBR increases observed after PMv-M1 ccPAS, reversed order ccPAS led to moderate PMBR reductions. Although there are strong projections from PMv to M1, projections from M1 to PMv also exist [29]. The moderate decrease of PMBR after M1-PMv ccPAS may, therefore, reflect not just a reduction in influence exerted by PMv over M1 but a change in the projections in the opposite direction.

Theta oscillations have been suggested as spectral fingerprints of top-down executive control [15-18]. Here we observed increased theta oscillations in No-Go trials after PMv-M1 ccPAS suggesting greater top-down motor control during response conflict as a result of entrainment of PMv-M1 connections. Opposite effects on theta oscillations are observed after reversed order M1-PMv ccPAS suggesting decreased executive control over motor output. Notably, while the ccPAS may cause some changes in early evoked and later induced theta activity (Fig.4) these modulations cannot be explained by changes in phase-locked responses (Fig.S3) or in ERP components (Fig.S4). Instead, the ccPAS appears to affect the amplitude of oscillatory activity linked to response inhibition [17, 32]. The results are also consistent with previous investigations emphasizing theta oscillatory activity in integrative mechanisms and as mediators of information transfer between prefrontal and motor areas in decision making and action control [33, 34].

Given the clear influence of ccPAS on theta oscillations during action inhibition in Experiment 3, changes in M1 cortical excitability as measured by MEP amplitude might, therefore, also have been expected in No-Go trials after ccPAS in Experiments 1+2. In fact, they were not seen. However, note that ccPAS effects on beta and theta oscillations manifested at specific timings after stimulus onset 700ms and 150ms after Go and No-Go stimuli respectively. By contrast, following standardized protocols, in Experiments 1+2, the PMv conditioning pulses in Baseline and Expression were delivered earlier, 125ms after on both Go and No-Go stimulus onsets. Therefore, it is possible that lack of modulation in M1 cortical excitability on No-Go trials in Experiments 1+2 may simply reflect the difficulty of probing PMv-M1 interactions across many time periods when using MEPs as indices of circuit activity.

In summary, cortico-cortical communication frequencies in the human PMv-M1 pathway can be manipulated in a state dependent manner. The frequency-specific patterns of oscillatory activity change found after different types of ccPAS on Go *vs* No-Go trials reflects spectral fingerprints of augmentation *vs* reduction of top-down PMv influence over M1. The patterns are consistent with Hebbian-like [4] spike-timing dependent long-term potentiation and depression and also consistent with hierarchical models of action control in which top-down motor control occurs in tandem with bursts of oscillations with specific resonant properties in the beta- and theta frequency ranges related to action termination and motor conflict and inhibition respectively.

## Acknowledgments

Funders: Bial Foundation (Grant 44/16), Templeton Grant, MFSR (Wellcome Trust: WT100973AIA) Data collection: Nadescha Trudel

## Author Contributions

Conceived and designed the experiments: AS MR. Performed the experiments: AS KA RD. Analyzed the data: AS KA RD LV MKF. Wrote the paper: AS MR LV MKF.

## Lead contact

Further information and requests for resources and reagents should be directed to and will be fulfilled by the Lead Contact, Alejandra Sel

