## Supplemental Information for "Entraining corticocortical plasticity changes oscillatory activity in action control and inhibition"

### Supplemental Figures

**Figure S1**

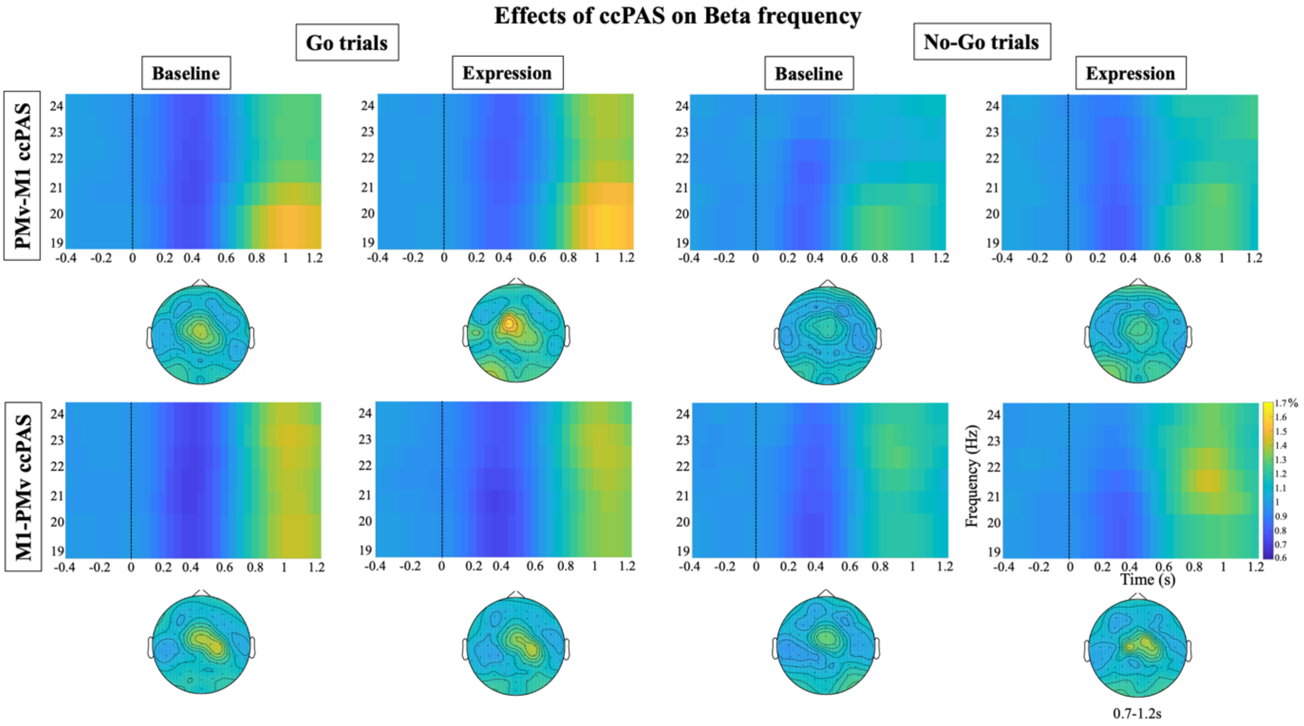


**Figure S1:** EEG time-frequency responses in the beta band (19-24Hz) in fronto-central sites (negative cluster; C4, CZ, FC2, CP2, FCZ, C1, C2, FC4, CP4, CPz) time-locked to the onset of the Go/No-Go stimuli, separated for Go and No-Go trials, and Baseline and Expression blocks, for Experiment 3 (top row) and Experiment 4 (bottom row).

### Figure S2

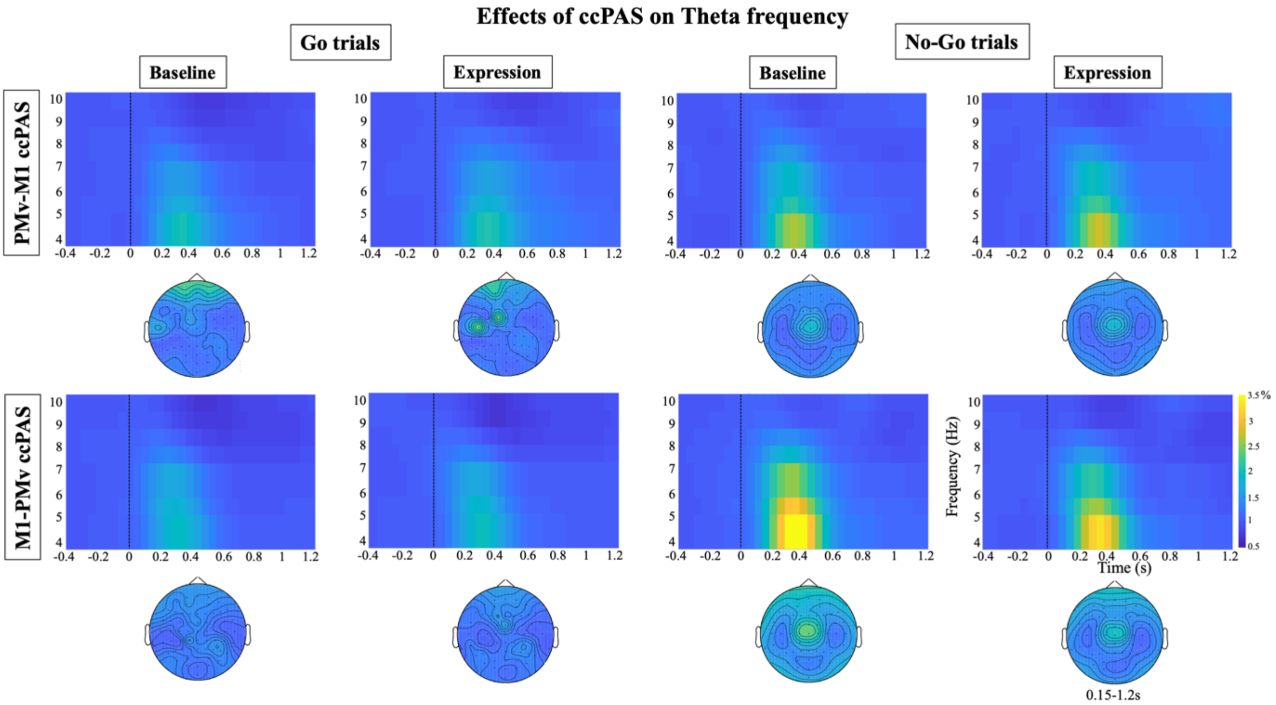


**Figure S2:** EEG time-frequency responses in the theta band (4-10Hz) in fronto-central sites (positive cluster; C3, C4, CZ, FC1, FC2, FCZ, C1, C2, FC3, FC4, CP4, CPZ) time-locked to the onset of the Go/No-Go stimuli, separated for Go and No-Go trials, and Baseline and Expression blocks, for Experiment 3 (top row) and Experiment 4 (bottom row).

**Figure S3**

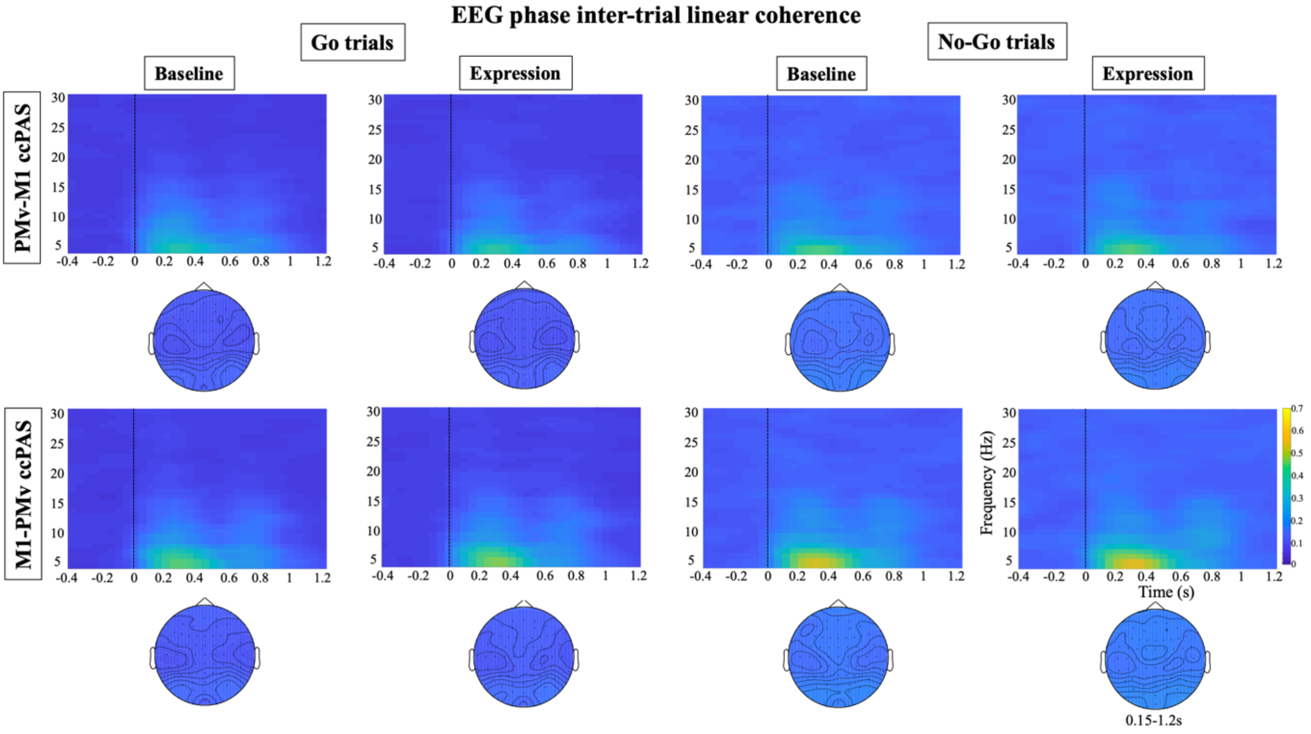


**Figure S3:** EEG phase inter-trial linear coherence in all bands tested (4-30Hz) in the main ROI (FC3, FC1, FCZ, FC2, FC4, C3, C1, CZ, C2, C4, CP3, CP1, CPZ, CP2, CP4) time-locked to the onset of the Go/No-Go stimuli, separated for Go and No-Go trials, and Baseline and Expression blocks, for Experiment 3 (top row) and Experiment 4 (bottom row). The results of the cluster-based permutation analysis on phase inter-trial linear coherence show a main effect of trial type ($p=0.001$) between 0.15 and 1.2s after stimulus onset for all frequencies tested 4-30Hz, indicating a greater coherence for No-Go trials *versus* Go trials in both Baseline and Expression blocks, irrespectively of ccPAS stimulation order. No other main effects or interaction were significant ($p>0.05$), indicating that the ccPAS protocol did not have an effect on phase coherence.

**Figure S4**

**ERPs to Go and No-Go trials**

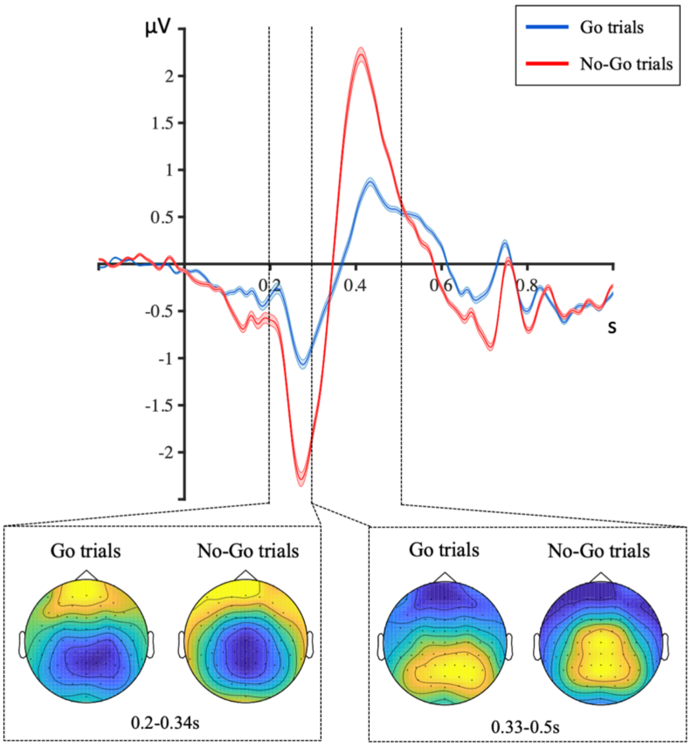


**Figure S4:** Visual ERPs time locked to the onset of the Go (blue line) and No-Go (red line) stimuli (averaged across electrodes - C3, C4, CZ, FC1, FC2, CP1, CP2, FCZ, C1, C2, FC3, FC4, CP3, CPZ), collapsed across the Baseline and the Expression block, and across Experiments 3 and 4 – shaded lines represent standard error for each condition. Topo-plots illustrate ERP activity in the time windows where the main effect of trial type was found. The results of the cluster based premutation analysis on visual ERPs revealed a significant difference between Go vs No-Go trials at 0.20 - 0.34s (positive cluster; p = 0.001; electrode sites - C3, CZ, FC1, FC2, CP1, CP2, FCZ, C1, C2, FC3, CP3, CP4, CPZ), and 0.33 – 0.5s after stimulus onset (negative cluster; p= 0.001; electrode sites - C3, C4, CZ, FC1, FC2, CP1, CP2, FCZ, C1, C2, FC3, FC4, CP3, CPZ), coinciding with the latency and distribution of the N2 and P3 component. No other main effects or interaction were significant (p>0.05), indicating that the ccPAS protocol did not have an effect on visual ERPs.

**Supplemental tables**

**Table S1**

**Accuracy for Go/No-Go trials in Baseline & Expression blocks**

| | | Go trials | No-Go trials |
|---|---|---|---|
| Experiment 1 | Baseline block | 0.95 ± 0.02 | 0.93±0.05 |
| | Expression block | 0.94 ± 0.02 | 0.92 ± 0.08 |
| Experiment 2 | Baseline block | 0.92 ± 0.06 | 0.94 ± 0.06 |
| | Expression block | 0.94 ± 0.02 | 0.89 ± 0.09 |
| Experiment 3 | Baseline block | 0.96 ± 0.02 | 0.89 ± 0.11 |
| | Expression block | 0.93 ± 0.02 | 0.87 ± 0.11 |
| Experiment 4 | Baseline block | 0.95 ± 0.01 | 0.90 ± 0.07 |
| | Expression block | 0.96 ± 0.01 | 0.89 ± 0.08 |

mean ± SD

**Table S2**

**Reaction time for Go/No-Go trials in Baseline & Expression blocks**

| | | Go trials |
|---|---|---|
| Experiment 1 | Baseline block | 393.38 ± 9.93 |
| | Expression block | 385.02 ± 8.04 |
| Experiment 2 | Baseline block | 399.67 ± 12.13 |
| | Expression block | 384.67 ± 8.95 |
| Experiment 3 | Baseline block | 360.38 ± 28.21 |
| | Expression block | 366.61 ± 38.06 |
| Experiment 4 | Baseline block | 345.02 ± 22.28 |
| | Expression block | 337.44 ± 18.64 |

median ± SD

### Supplemental Experimental Procedures

#### Participants

Eighty-three healthy, right-handed adults participated across four experiments. Twelve participants were excluded due to equipment failure and three participants were excluded due to excessive noise in the EEG signal resulting in 68 participants – 18 in Experiment 1 (24 ± 7.13; 10; 0.83 ± 0.13); 17 in Experiment 2 (23.94 ± 4.07; 7; 0.90 ± 0.09); 16 in Experiment 3 (23.75 ± 4.59; 10; 0.81 ± 0.17); 17 in Experiment 4 (22.64 ± 2.31; 5; 0.93 ± 0.13) (where numbers correspond to mean age ± SD; number of female participants, handiness mean ± SD; as measured by the Edinburgh handedness inventory- adapted from [S1]). Some subjects participated in more than one experimental session for Experiments 1+2, but in this case, sessions were on average, 32.91 ± 22.58 days (mean ± SD) apart. The number of naïve and non-naïve participants were similar across both experiments. All participants had no personal or familial history of neurological or psychiatric disease, were right handed (except for one participant – handiness score .045), gave written informed consent (Medical Science Interdivisional Research Ethics Committee, Oxford RECC, No. R29477/RE004), were screened for adverse reactions to TMS and risk factors by means of a safety questionnaire, and received monetary compensation for their participation. Participants underwent high-resolution, T1-weighted structural MRI scans. Sample sizes were determined based on previous studies that have used the same ccPAS protocol to measure the influence of PMv over M1 cortical excitability [S2, S3](Experiments 1+2), and studies that have used the Go/No-Go paradigm to investigate oscillatory responses during action control in humans (Experiments 3+4)

#### Experimental design

All four experiments started with a Baseline block, followed by a ccPAS period, and an Expression block (Fig.1). During Baseline and Expression blocks participants performed a visual Go/No-Go task. Trials started with the presentation of either a blue (Go trials – 70% of trials) or a red (No-Go trials) square (1.8 x 1.8cm) displayed for 1s, or until response, in Experiments 1+2, and for 500ms in Experiments 3+4. These were followed by a yellow fixation cross (1.3 x 1.3cm) presented centrally on the screen for a time interval between 0.5s and 3.8s in Experiments 1+2, and between 2s and 3s in Experiments 3+4. There was a total of 468 trials per block in Experiments 1+2, and 304 trials per block in Experiments 3+4 (equal number of trials in the Baseline and the Expression blocks), with a short break half way through the block. Blocks always started with four consecutive go trials. Participants were instructed to press a button with their left index finger as soon as the blue square was presented, and to withhold the response when the red square appeared on the screen. Reaction times and accuracy were recorded. During the task, participants were seated at approximately 50cm from the screen in an air-conditioned and sound-attenuated room in Experiments 1+2, while in Experiments 3+4 they sat in a sound and electrically shielded booth.

In Experiments 1 and 2, TMS over PMv and M1 was delivered while participants performed the task during the Baseline and Expression blocks. The effect exerted by PMv stimulation on subsequent M1 stimulation was determined by contrasting motor-evoked potentials (MEPs) recorded from left first dorsal interosseus (FDI) muscle, when either single-pulse TMS (spTMS; 72 trials per block) was applied to right M1, or paired-pulse TMS (ppTMS) was delivered over both right PMv and M1 (72 trials per block). Both resting-state and task-state interactions between M1 and PMv, and adjacent areas, emerge at 6–8 ms intervals [S2-S5]. Precise interpulse timing is critical if both PMv and M1 pulses are to produce coincident influences on corticospinal activity. Therefore, we applied an interpulse interval (IPI) of 8 ms. The pulse applied to M1 always occurred 125ms after the stimulus onset. This stimulation time was chosen based on a pilot study (N=10) showing a greater influence of the state-dependent (Go vs No-Go) influence of PMv on M1 in ppTMS trials at 125ms (in contrast to 100, 150, 175ms), and it accords with the early engagement of motor and pre-motor areas in action control shown in previous studies [S6, S7]. Both spTMS and ppTMS trials were administered in alternation with no-TMS trials (324 trials per block). Trial presentation order was pseudorandomised within the same block of trials to ensure that there were no more than five consecutive TMS trials to avoid TMS after-effects.

In all four experiments, the ccPAS period that intervened between Baseline and Expression blocks consisted of 15 min of ccPAS over PMv and M1 applied at 0.1 Hz (90 total stimulus pairings) with an IPI of either 6 or 8 ms[1]. In Experiments 1 and 3, the pulse applied to PMv always preceded the pulse over M1, while the opposite was true in Experiments 2 and 4, which served as an active control.

[1] Due to error in the experimental set up, the IPI was 6ms for half of the participants in Experiment 3 and for half of the participants in Experiment 4. The impact of this variance in the experimental manipulation was tested by a repeated measures ANOVA with within-subject factors block (Baseline, Expression) and trial type (Go, No-Go), and between-subject factors ccPAS order (forward, reverse), and IPI (8ms, 6ms). No effects of the 6ms IPI versus 8ms IPI was seen even when the analysis focused on the time window and frequency bands in which

**TMS and electromyography recordings**

TMS was applied using two Magstim 200 stimulators, each connected to 50 mm figure-eight coils. The M1 "scalp hotspot" was the scalp location where the TMS stimulation evoked the largest left FDI MEP amplitude. This scalp location was projected onto high-resolution, T1-weighted MRIs of each volunteer's brain using frameless stereotactic neuronavigation (Brainsight; Rogue Research). In contrast to the scalp hotspot, the right M1 "cortical hotspot" was the mean location in the cortex where the stimulation reached the brain for all participants in Montreal Neurological Institute (MNI) coordinates ($X = 41.37 \pm 7.05$, $Y = -14.74 \pm 9.01$, $Z = 64.71 \pm 7.35$; Fig. 1) was similar to that reported previously [S2, S3, S5, S6]. The PMv coil location was determined anatomically as follows. A marker was placed on each individual's MRI and adjusted with respect to individual sulcal landmarks to a location immediately anterior to the inferior precentral sulcus. The mean MNI cerebral location of the PMv stimulation was at ($X = 58.78 \pm 3.13$, $Y = 18.03 \pm 6.96$, $Z = 16.13 \pm 10.77$; Fig 1) and lies within the region defined previously as human PMv [S8], more precisely over areas 44d and 44v of the pars triangularis within the inferior frontal gyrus [S9], which resembles parts of macaque PMv in cytoarchitecture and connections [S10, S11].

Resting motor threshold (RMT) of the right M1 (mean ± SD, 43.13 ± 7.22% stimulator output) was determined as described previously [S12]. As in previous ccPAS studies [S2, S3, S6, S13], PMv TMS was proportional to RMT - 110% (47.76 ± 7.35). M1 stimulation intensity during experiments was set to elicit single-pulse MEPs of ±1 mV (47.23 ± 7.58% stimulator output). TMS coils were positioned tangential to the skull, with the M1 coil angled at ~45° (handle pointing posteriorly), and the PMv coil at ~0° relative to the midline (handle pointing anteriorly). The PMv coil was fixed in place with an adjustable metal arm and monitored throughout the experiment. The M1 coil was held by the experimenter.

Left FDI electromyography (EMG) activity was recorded with bipolar surface Ag-AgCl electrode montages. Responses were bandpass filtered between 10 and 1000 Hz, with additional hardwired 50 Hz notch filtering (CED Humbug), sampled at 5000 Hz, and recorded using a CED D440-4 amplifier, a CED micro1401 Mk.II A/D converter, and PC running Spike2 (Cambridge Electronic Design). Motor evoked potentials (MEPs) were computed as the peak-to-peak amplitude difference. Trials in which MEPs were under 0.1mV (trials in which no MEP was elicited, mostly because of coil misplacement or missing stimulation due to, e.g., coil overheating) or over 6mV were discarded.

As the distribution of MEP amplitudes was positively skewed, they were log-transformed for further statistical analyses. A repeated-measures analysis of variance (ANOVA) using the factors of block (Baseline, Expression), trial type (Go, No-Go) and TMS (spTMS, ppTMS), was used to analyse the electromyographic data in Experiments 1+2. To correct for non-sphericity, Greenhouse-Geisser corrected results are reported. Following previous observations showing greater MEP amplitudes when M1 TMS was preceded by a previous conditioning pulse over PMv in movement trials [S2-S5, S7, S13], we performed an additional analysis directly contrasting MEP amplitudes in spTMS *versus* ppTMS in Go trials, in the 12 subjects that participated in both Experiments 1+2.

**EEG recording and analysis**

EEG was recorded with sintered Ag/AgCl electrodes from 64 scalp electrodes mounted equidistantly on an elastic electrode cap (64Ch-Standard-BrainCap for TMS with Multitrodes; EasyCap). All electrodes were referenced to the right mastoid and re-referenced to the average reference off-line. Continuous EEG was recorded using NuAmps digital amplifiers (Neuroscan, El Paso, Texas 1000Hz sampling rate).

Off-line EEG analysis was performed using Fieldtrip [S14]. The data was down-sampled to 500Hz and digitally band-pass-filtered between 1-40 Hz. Bad/missing channels were restored using a FieldTrip based spline interpolation. Next, the data were segmented into 3.5s intervals starting from 1.4s before stimulus onset. This was done for Go and No-Go trials separately, and incorrect trials and trials where RTs were too slow (+/- 2SD) were excluded from the analysis. Automatic artefact rejection was combined with visual inspection for all participants eliminating large technical and movement related artefacts. Physiological artefacts such as eye blinks and saccades were corrected by means of independent component analysis (RUNICA, logistic Infomax algorithm) as implemented in the FieldTrip toolbox. Those independent components (most often one or two) whose timing and topography resembled the characteristics of the physiological artefacts were removed. The signal was re-referenced to the arithmetic average of all electrodes, and segments were baseline-corrected using an interval from - 500ms to - 100ms before the stimulus onset.

For the time-frequency analysis, single-subject activations for each block (Baseline, Expression) and trial type (Go, No-Go) were averaged and submitted to a complex multitaper time-frequency transformation from 4 to 30Hz in steps of 1Hz, with a fixed Hanning window of 0.75s. A relative Baseline normalization was

---

the key effects of ccPAS on neural oscillations had been found. The analysis merely confirmed the differential effects of ccPAS on beta and theta bands for Go and No-Go trials respectively.

performed using a time window from -1.1s to 0s in respect to stimulus onset. To estimate the effects of the ccPAS protocol on neural responses of action control in the Go/No-Go task, time-frequency activations time-locked to stimulus onset were computed at the group level using a non-parametric randomisation test controlling for multiple-comparisons [S15]. Investigations of the neural dynamics of cognitive and motor control processes highlight the functional significance of both low and high frequency oscillations in action performance and inhibition. Theta (4-8 Hz), alpha (9-12 Hz) and beta (13-30 Hz) spectrums have all been linked to aspects of action control. Therefore, in the statistical analyses no frequency bands were selected *a-priori*. Instead, the statistical analyses were performed on all motor relevant frequency bands (4-30Hz) and across the entire time window where oscillatory changes associated with motor control have been noted to take place – this is, 0.2-1.2s after stimulus onset. Statistical analyses were restricted to 15 electrodes distributed over frontocentral and parietal areas, i.e. FC3, FC1, FCZ, FC2, FC4, C3, C1, CZ, C2, C4, CP3, CP1, CPZ, CP2, CP4, where the neural phenomena linked to motor control are typically distributed [S16-S18].

To test if the ccPAS protocol influenced cortical correlates of action control, and if this influence happened in a state-dependent manner (Go vs. No-Go), we first computed the 'cortical entrained effect' (calculated by subtraction of each frequency at each time point of activity recorded in Baseline from the Expression block) for Go and No-Go trials separately. We then calculated the difference of the cortical entrained effect between No-Go trials vs Go trials. Thereafter, we contrasted the 'No-Go-minus-Go cortical entrained effect' recorded from the participants that received PMv-M1 ccPAS (Experiment 3; N=16), vs. the participants that received reverse order M1- PMv ccPAS (Experiment 4; N=17), by means of between subject non-parametric cluster-based permutation analysis. A non-parametric, cluster-based permutation approach is an efficient way of dealing with the multiple comparison problem that prevents biases in selecting time-windows or electrode sites avoiding inflation of type I error rate [S15, S19]. Time frequency responses in all conditions are represented in Figure S1(beta band) and Figure S2 (theta band).

Subject-wise time-frequency courses were extracted at the selected electrodes and were passed to the statistical analysis procedure in FieldTrip, the details of which are described by Maris and Oostenveld [S15]. Subject-wise time-frequency courses were compared to identify statistically significant clusters in the time, frequency, and spatial domain using a FieldTrip-based analysis [S14]. FieldTrip uses a nonparametric method [S20] to address the multiple comparison problem. T-values of adjacent temporal and frequency points whose p-values were less than 0.05 were clustered by adding their t-values, and this cumulative statistic is used for inferential statistics at the cluster level. This procedure, i.e. the calculation of t-values at each temporal point followed by clustering of adjacent t-values, was repeated 5000 times, with randomised swapping and resampling of the subject-wise time-frequency activity before each repetition. This Monte Carlo method results in a nonparametric estimate of the P-value representing the statistical significance of the identified cluster.

In addition, to rule out the possibility that changes in oscillatory activity after ccPAS were linked to phase-locked responses to stimulus presentation, we computed the phase coherence across trials (inter-trial linear coherence - ITLC) for each condition (Figure S3). We tested the effects of ccPAS on ITLC mimicking the cluster-based permutation analysis performed on time-frequency oscillatory responses across all time points and frequency bands.

Visual observation of the No-Go trials in the group undergoing reverse order M1-PMv ccPAS revealed a particularly high theta power in three participants. To ensure that the effects observed in the theta range were not limited to participants with especially high or low theta power we performed a control analysis in which we left out the three participants with higher theta scores and the three participants with lower theta scores in the window of interest (0.15 – 1.2s after stimuli onset). We repeated the cluster-based permutation analysis to test differences in the cortical entrained effect in No-Go trials in the theta band and we observed that the significant differences in No-Go trials between PMv-M1 ccPAS *versus* M1-PMv ccPAS remained (positive cluster: p =0.005; electrode sites - CZ, FC1, FC2, FCZ, C1, C2, FC3, FC4, CPZ; between 0.15 and 0.55s after stimulus onset), indicating that this interaction was not driven by the group differences in theta power at baseline.

For the event related potential (ERP) analysis, single-subject ERPs for each block (Baseline, Expression) and trial type (Go, No-Go) were calculated and used to compute ERP grand averages across subjects (Figure S4). The analysis on the ERP data mimicked the time-frequency analysis. In brief, ERP activations time-locked to stimulus onset were computed at the group level using a non-parametric randomisation test controlling for multiple-comparisons [S15]. To test the effects of ccPAS on ERPs related to action control, we first computed the ccPAS effect on ERPs (by subtraction of each time point of the trials in the Baseline block from the Expression block) for Go and No-Go trials. We then computed the difference of the ccPAS effect between No-Go and Go trials. Finally, we contrasted the 'No-Go-minus-Go ccPAS effect' between the two participant groups (PMv-M1 ccPAS group vs reversed order PMv-M1 ccPAS group) by means of between subject non-parametric cluster-based permutation analysis. Statistical analyses were done across the entire time window where the N2-P3 component typically takes place, this is, 0.2-0.6s [S21] and it was restricted to 15 electrodes distributed over frontocentral and parietal areas (see above). Subject-wise activation time courses were extracted at the selected electrodes and were passed to the analysis procedure of FieldTrip

[S15]. The cluster-based permutation analysis on the ERP data did not find any significant differences in the cortical entrained effect between the participant groups in Experiments 3 and 4 at any electrode cluster when contrasting either Go or No-Go trials. These results demonstrated that (1) the effects of rPAS on the PMv-M1 circuit are frequency specific and only affect particular oscillatory bands linked to action control, i.e. beta and theta bands; (2) the changes observed in the slow-frequency band theta cannot be explained by changes in the ERP components. There was, however, a significant difference between Go *versus* No-Go trials across both groups, confirming that the action control manipulation was effective.

**Behavioural analysis**

Behavioural performance measures comprised median RTs (excluding trials with RT +/- 2SD from the mean) and accuracy (excluding omission errors in go trials and commission errors to No-Go trials). We tested the effect of the ccPAS protocol on RTs and accuracy measures. In Experiments 1+2, analyses focused on data from no-TMS trials to avoid bias from the TMS impact on the hand muscles controlling the button press [S22]. A repeated-measures analysis of variance (ANOVA) using the within-subject factors of block (Baseline, Expression) and trial type (Go, No-Go) was used to analyse the behavioural data of Experiments 1+2 – i.e. two separate ANOVAs, one for each Experiment. For Experiments 3+4, these analyses also included the between-subject factor of ccPAS order (PMv-M1 or reversed order M1-PMv), and within-subject factors of block (Baseline, Expression), and trial type (Go, No-Go). The behavioural analysis did not show significant differences in accuracy or median RT between Baseline and Expression blocks in any of the 4 experiments ($ps > 0.05$). (Supplemental Table 1 and 2).

In Experiments 3+4, we explored the possibility that MEP/EEG modulations (computed as the difference between Baseline and Expression blocks) could be linked to participants' performances (median RT and accuracy rates) at Baseline. No relationship between was found participants' median RT/accuracy in Go trials and MEP changes between the Baseline vs. Expression blocks in Go trials in Experiment 1 ($p > 0.05$). We also tested if undergoing ccPAS influenced the after-effects of No-Go trials on subsequent Go trials, and found that there were aftereffects of No-Go trials on subsequent Go trials represented by slower median RTs in the Expression *vs* Baseline period ($F(1,31) = 7.746$, $p = 0.009$, $\eta p^2 = 0.2$), possibly due to fatigue.
